# DAIRYdb: A manually curated gold standard reference database for improved taxonomy annotation of 16S rRNA gene sequences from dairy products

**DOI:** 10.1101/386151

**Authors:** Marco Meola, Etienne Rifa, Noam Shani, Céline Delbès, Hélène Berthoud, Christophe Chassard

## Abstract

Reads assignment to taxonomic units is a key step in microbiome analysis pipelines. To date, accurate taxonomy annotation, particularly at species rank, is still challenging due to the short size of read sequences and differently curated classification databases. However, the close phylogenetic relationship between species encountered in dairy products requires accurate species annotation to achieve sufficient phylogenetic resolution for further downstream ecological studies or for food diagnostics. Taxonomy annotation in universal 16S databases with environmental sequences like Silva, RDP or Greengenes is based on predictions rather than on studies of type strains or isolates. We provide a manually curated database composed of 10’290 full-length 16S rRNA gene sequences from prokaryotes tailored for dairy products analysis (https://github.com/marcomeola/DAIRYdb). The performance of the DAIRYdb was compared with the universal databases Silva, LTP, RDP and Greengenes. The DAIRYdb significantly outperformed all other databases independently of the classification algorithm by enabling higher accurate taxonomy annotation down to the species rank. The DAIRYdb accurately annotates over 90% of the sequences of either single or paired hypervariable regions automatically.

The manually curated DAIRYdb strongly improves taxonomic classification accuracy for microbiome studies in dairy environments. The DAIRYdb is a practical solution that enables automatization of this key step, thus facilitating the routine application of NGS microbiome analyses for microbial ecology studies and diagnostics in dairy products.

## Introduction

The exploration of microbial communities has experienced a boost during the last decade with the advent of next generation sequencing (NGS) technologies (1). Previously undetectable, since unculturable, micro-organisms in soils (2), water (3, 4), airborne (5, 6), snow (7), ice (8), food (9), human gut (10–12) etc. could be unravelled at an unprecedented depth and resolution. An infinite number of descriptive studies have been published describing microbial community structures in various environments, often correlating their dynamic changes over time or space by means of the 16S rRNA gene (13–15).

### The marker gene 16S rRNA

The 16S rRNA gene (16S) was proposed by Carl Woese and George Fox in the 1980s as the gold standard marker gene for molecular taxonomic research (16–18). Several characteristics of the 16S make it a marker gene for surveys of microbial diversity: i) it is an ubiquitous and highly conserved gene in prokaryotes, ii) it has a functional degree and size presenting clock-like mutation rates like an evolutionary chronometer, iii) and the presence of alternating conserved and hypervariable regions (HVRs) permit to design universal primers on the conserved regions and to use the HVRs (V1-V9) for taxonomic classification (19, 20).

Before the advent of NGS, microbial community studies were based on fingerprinting techniques, such as DGGE, TRFLP or LH-PCR sometimes in combination with Sanger sequencing of the complete 16S spanning over about 1550 bp. While Sanger sequencing delivered almost the complete 16S at good quality, the throughput was low due to the high workload preventing researchers to unravel the full array of microbial diversity within a sample (20).

### Classification tools

Taxonomic classification of the 16S is not trivial and requires both familiarity with prokaryotic phylogeny and often manual intervention due to poor annotation of the operational taxonomic units (OTUs) by the available 16S databases (21). On the one hand, NGS has triggered the acquisition of enormous amounts of sequencing data, offering the possibility to overcome the limitations of Sanger sequencing. On the other hand it has brought huge computational challenges, such as the risk of taxonomic missannotations or ambiguous results during taxonomic classification steps. Despite the steadily increasing read length obtained by NGS, the need for trustworthy classification of very short 16S sequences covering only one to three HVR remains a crucial step to obtain robust and accurate taxonomic classification in modern microbiology (22).

Numerous classification predictors algorithms have been developed and optimized in recent years with the aim to accurately annotate the taxonomy of OTUs from short reads. Those classification tools have been developed for 16S and other genes based on different mathematical models, such as *e.g.*, k-mer, Bayesian, Hidden Markov-Monte-Carlo model (HMM) etc.). The Basic Local Alignment Search Tool (Blast) has long been the gold standard for sequence comparison and annotation (23). In recent years, more 16S specific taxonomy predictors have been developed, including RDP Naive Bayesian Classifier (NBC) (24), a naive Bayesian Classifier based on k-mers, GAST (25), MEGAN (26), Metaxa2 (27), riboFrama (28), SPINGO (29), PROTAX (30), SINTAX (31), DynamiC (32), Humidor (33), MAPseq (34), microclass (35) and other tools implemented in the most current 16S pipelines like mothur (36), Qiime v1 (37), Qiime v2 (https://qiime2.org) and FROGS (38).

Here we used three taxonomy predictors based on different algorithms and programming languages (39), in which dedicated databases can be integrated and used for classification. One of the three classification tools used in this study to test the performance of the DAIRYdb was the updated version of Blast, Blast+ (40). It is based on an heuristic method for identification of database sequences that resemble the query sequence above a certain threshold. It searches for short sequence matches, which are locally aligned after the first match (40).

Another classification tool used in this study was Metaxa2, an HMM based software tool for automated detection and classification of short NGS fragments, such as ribosomal small and large subunits, SSU and LSU, respectively, or any gene of interest useful for classification of any organism (27, 41). The third taxonomy predictor used in this study was SINTAX, a non-Bayesian taxonomy classifier specific for 16S sequences, which uses k-mer similarity to identify the top hit in a reference database providing bootstrap confidence values at each taxonomic rank (31).

### 16S repositories

Although classification prediction algorithms have strongly improved, manually curated databases containing only authoritative full-length 16S sequences from type strains and cultivated reference strains can potentially compensate the limitations of short read sequences annotations by means of sophisticated algorithms. To date, three main independent universal repositories dedicated to universal 16S sequences from prokaryotes are widely used: Silva, The Ribosomal Database Project (RDP), and Greengenes (42).

Silva is the universal 16S repository with the highest number of sequences. The latest release of Silva SSU/LSU 132 (www.arb-silva.de) contained 6’073’181 16S sequences of at least 300 bp, with 2’090’668 good quality sequences with at least 900 bp length (43–45). Taxonomic rank information of Silva and Living Tree Project (LTP) are based on the Bergey’s Taxonomic Outlines and the List of Prokaryotic Names with Standing Nomenclature (LPSN) (46). Minimal training sets, such as the SSU Ref NR 99 or the LTP (47), offer a reduced number of sequences for faster classification but still covering the broadest currently known biodiversity.

The second biggest repository, the Ribosomal Database Project (RDP Release 11, Update 5; http://rdp.cme.msu.edu) (48), contained at the time of writing 3’356’809 16S sequences from the International Nucleotide Sequence Database Collaboration (INSDC) (49). The nomenclature is based on the Bacterial Nomenclature Up-to-Date and the taxonomic rank information on the Bergey’s Manual.

Greengenes v13_5 (50) contains 1’800’000 quality filtered 16S sequences. Classification nomenclature is based on automatic *de novo* tree construction and rank mapping with the NCBI Taxonomy database (51). Although frequently used in community studies together with Qiime (37), the last update dates back to 2013 with no indication for an imminent update.

### Taxonomy annotation in microbiology

Phylogenetic classification has tailored taxa by means of phylogenetic, phenotype and genomic coherence that make taxonomic units unique within the classification schema (52). Phylogenetic coherence is determined by the 16S for which the previously mentioned databases provide a valuable tool for taxonomic classification (52). However, the exponential increase of 16S sequences from previously unknown and uncultured bacteria led to an explosion of exotic labels at any taxonomic rank with often contrasting taxonomic classifications between databases based on different taxonomic catalogues (*e.g.*, Bergey’s Taxonomic Outlines, List of Prokaryotic Names with Standing Nomenclature (LPSN), International Sequence Database Collaboration (INSDC), Bacterial Nomenclature Up-to-Date (53, 54)).

The lack of consensus on a widely accepted taxonomy, as well as the lack of taxonomic characterisation of yet uncultured bacteria, are severely limiting communication among scientists and may lead to incorrect annotations and thus "poison every experiment that makes use of them" (52, 55). In worst cases, incorrectly annotated bacteria are included in databases further used to classify new sequences. In fact, it has been shown that there are numerous unambiguous disagreements between the nomenclature hierarchies, where many taxon names are placed into different parent taxa in the databases Silva and Greengenes or the taxonomy is not consistent with the tree (54). In addition to hierarchy disagreements, about 34% of identical sequences in Silva and Greengenes databases presented annotation conflicts, 24% of which were blanks (unclassified) in either of both database (54). While the blanks imply a false negative by one database or a false positive by the other, differently annotated sequences with the same Accession number or identical sequence string must be due to annotation error in one or both databases (54).

While NGS has allowed researchers to obtain deep insights into the microbial community structures inhabiting various environments, the complexity of the analytical process and taxonomy annotation on short read sequences is still challenging and prevents researchers to deploy microbial community analysis for diagnostic purposes in an accurate and reproducible way (1, 67). Previous studies have highlighted the importance of high-quality data for improving the classifications of the obtained OTUs (22, 68, 69). Although universal 16S databases cover vast prokaryotic biodiversity, they often fail to guarantee accurate classification to the species rank for sequences obtained from a highly studied environment, such as dairy products. In fact, classification accuracy at lower taxonomic ranks increases with a gold standard training set encompassing only full-length and good quality representative sequences innate to the investigated environment (22, 54, 69, 70).

In microbiology, distinction is made between the concept and the definition of species (52). The species concept explains the idea of what is considered to be a species as a unit of biodiversity, the meaning of the patterns of recurrence observed in nature, and the reason for their existence (71). The species definition, however, is concretely the set of parameters that are applied to circumscribe the category (72).

Thanks to the dropping costs, NGS is increasingly applied routinely as diagnostic technology for quality assessments and microbial community analyses in dairy products. Several initiatives aimed at tracking from "Farm to Fork" the range of expected microbial communities along the food supply chain, such as Food Safety Consortium with IBM, Mars and Bio-Rad Laboratories (www-03.ibm.com/press/us/en/pressrelease/45938.wss and www-03.ibm.com/press/us/en/pressrelease/52690.wss/). However, fast and accurate, thus automatized classification of the OTUs is not yet possible at the biologically most significant species rank due to the short sequence fragments and the absence of food-dedicated, thoroughly curated 16S databases, particularly. Therefore, manually curated databases are of paramount importance to improve reproducibility, speed during the bioinformatics process of microbial community studies and communication between researchers (54).

Here we present a comprehensive gold standard database, DAIRYdb (Database, Agroscope, Inra, Ribosomal, accuracY), for 16S OTUs classification from NGS data of dairy products. The main goal was to develop a dedicated database that allow researchers to accurately and automatically annotate short reads of 16S down to the species level. Manual curation of the database and its restriction to the biodiversity expected in dairy products strongly improves accuracy and reproducibility of phylogenetic classification to all taxonomic ranks. DAIRYdb is publicly available at https://github.com/marcomeola/DAIRYdb and can be integrated in any classification prediction tool that allows the integration of customized databases, such as Blast+, Metaxa2, SINTAX and FROGS.

## Results

### Construction

The 16S sequence database of dairy products DAIRYdb was constructed using a set of over 390’000 sequences associated to the selected keywords (cheese, milk, teat, dairy, starter, whey) deposited in NCBI GenBank and ENA/EMBL, as well as sequences with 97% ANI from Silva, RDP and Greengenes (Figure 1). About 10’000 best quality reference sequences were retained after filtering based on sequence length (*>*1300 bp), quality (pintail *>*75) and potential chimeras. Finally, 16S sequences of important species from cheese and dairy environments (73, 74), whose sequences were lost during the clustering, were added, resulting to the final number of 10’290 16S sequences.

**Fig. 1.**
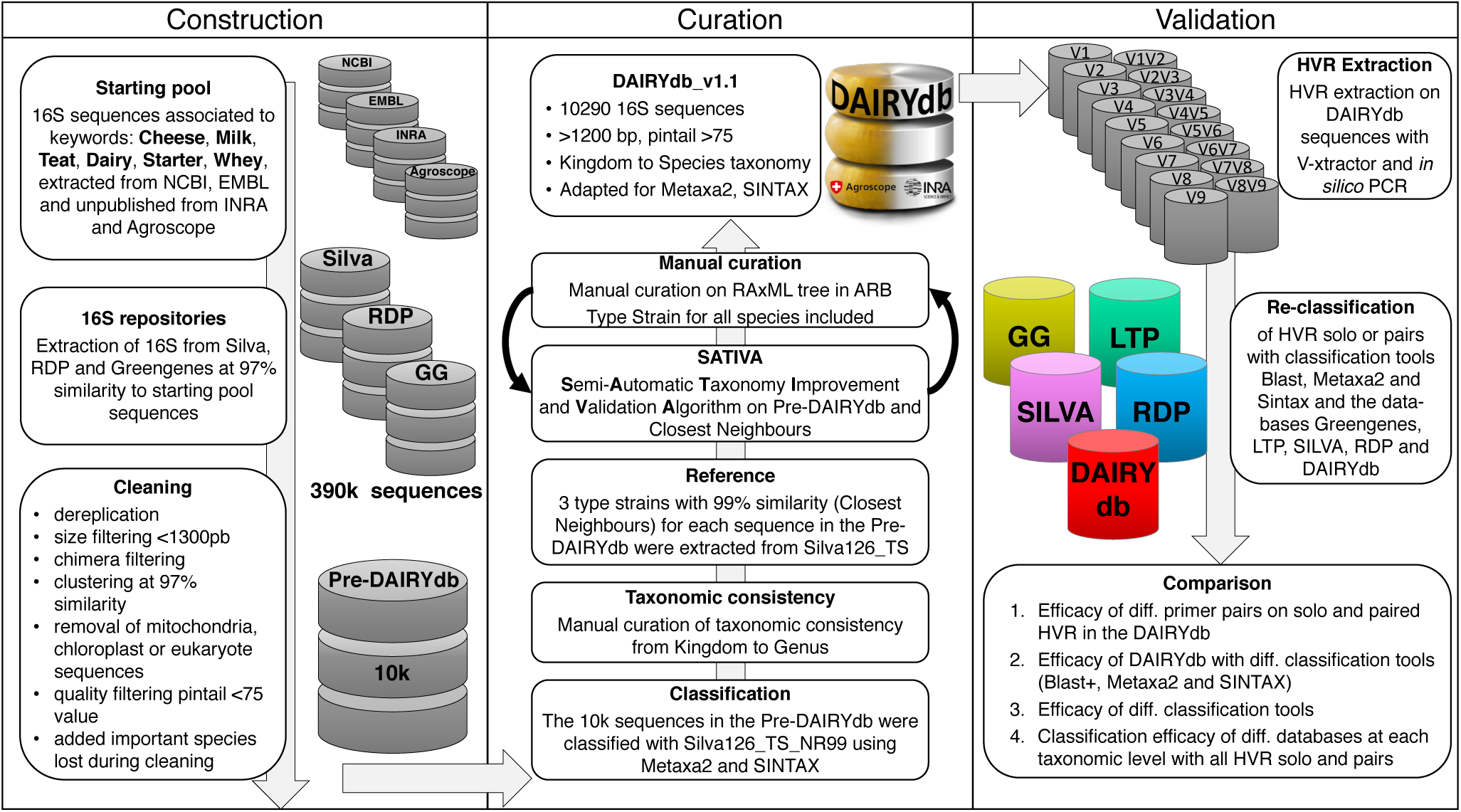
Development of the DAIRYdb consisted in three main steps: construction, curation and validation. For construction, dairy products specific 16S rRNA gene sequences were retrieved from Silva, RDP and Greengenes using Genbank NCBI, EMBL, Agroscope and INRA sequences. Curation was performed based on the cross-validation results from the leave-one-out test of SATIVA and highly iterated RAxML tree, followed by manual curation of taxonomic assignment and consistency throughout all taxonomic ranks, with a particular focus on singleton taxons with no reference sequence. Validation was performed comparing identification accuracy of single and HVR pairs by the five databases (Greengenes 13.8, LTP version, Silva 128 NR99, RDP version and DAIRYdb).

The observed distribution among the different key words might reflect the unequal distribution of microbiome studies predominantly performed on cheese, dairy and milk samples, as compared to teats and whey. About 1933 sequences of the DAIRYdb were shared among all keywords (Figure 2A) and 1778 were shared among the keywords dairy, cheese and milk. In fact, the majority of the sequences composing the DAIRYdb were linked to those three keywords (Figure 2B). Altogether, 1’700 sequences were associated to just one keyword, with most of the sequences shared by four keywords (Figure 2C).

**Fig. 2.**
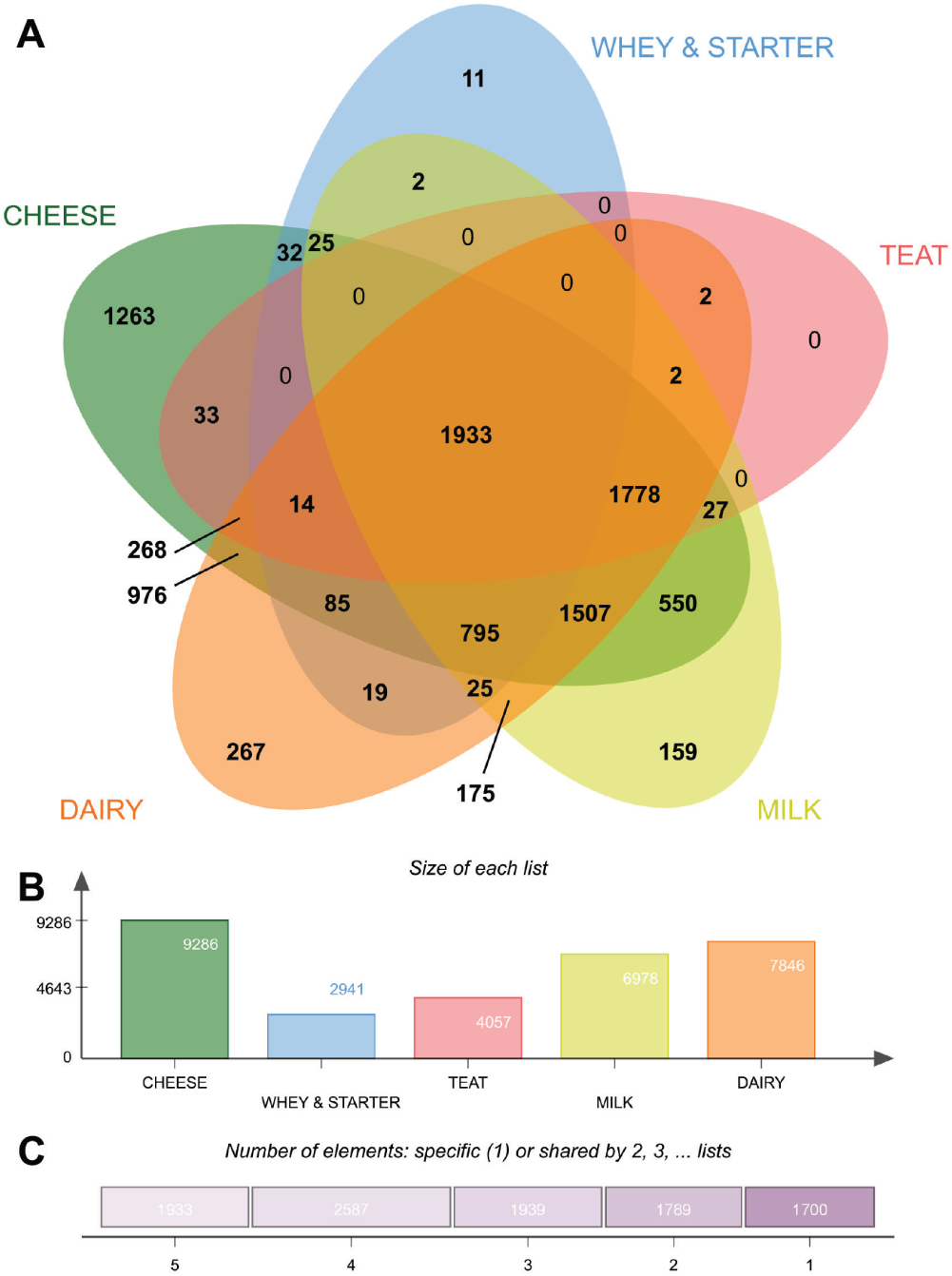
Origin of sequence in the DAIRYdb. A) Five-factors Venn diagram comparing the origins of the sequences (9’948) retrieved from the public repositories Genbank NCBI and EMBL associated to the keywords “cheese”, “dairy”, “milk”, “teat” and “whey/starter”. About 12.7% (1’263) sequences were only detected in cheese and 15.1% (1’507) were detected in all three cheese, milk and dairy environments. B) Total number of sequences associated to a particular keyword. C) Number of sequences shared by 1 to 5 keywords. About 19.4% (1’933) sequences were detected in all 5 keywords, while 17.1% (1’700) sequences were unique to one keyword.

### Curation

During the first step of data curation, the sequences were taxonomically annotated with Silva by means of SINA (75). The resulted annotation at all taxonomic ranks underwent a first manual check and cleaning for taxonomic inconsistencies through cross-comparison with the other members of the same taxonomic rank in a phylogenetic tree. No taxonomic overlaps comparable to other databases are present in the DAIRYdb, where different species of the same genus fall under different taxonomic lineages (54). A maximum of three closest neighbour type strains (CN) with authoritative taxonomy (CN) from Silva sharing 99% global sequence similarity to each sequence in the DAIRYdb were added to the 10’290 sequences in the DAIRYdb as reference during the curation process and removed at the end of the curation process. The maximal number of lowest common ancestors (LCA) with an authoritative taxonomy strongly improved the curation process with the Semi-Automatic Taxonomy Improvement and Validation Algorithm (SATIVA) increasing robustness of the proposed changes of miss-annotated environmental sequences within the DAIRYdb (76).

By using only near full-length and curated 16S from type strains as reference sequences, we were able to validate and correct the taxonomy annotation where necessary. The SATIVA results were inspected and taxonomy manually curated using a highly iterated phylogenetic tree. The approach used during the manual curation broadly follows the rationale described in detail in a recently published study (54). Taxonomy annotations from authoritative type strain sequences were used as reference for the environmental sequences in the tree. For ranks at which no taxonomic annotation was possible with certainty due to the lack of authoritative type strains within the same clade (*i.e.*, commonly labelled unknown, uncultured etc. in universal databases), the *lowest common rank* (LCR) (70) was used down to the species rank with the addition of the unclassified rank. Although not all OTUs in a microbiome study will be classified to the species rank, at least they will not all be merged to the same unclassified species rank, but taking over the LCR to differentiate between all unclassified OTUs. As an example, a sequence assigned to the LCR, the genus *Sporichthya*, was named at species rank *Sporichthya_Species*. This approach avoids the merging of abundance values from different unknown species to biological uninformative groups, thus improving communication among scientists (53).

DAIRYdb version 1.1 contains 2 kingdoms (Bacteria and Archaea), 47 phyla, 136 classes, 249 orders, 463 families, 1757 genera and 4030 unique species-like groups/species complexes (Figure 3, Additional File 1 and Additional File 2). The *Firmicutes* is the predominant phylum with 37% of all sequences, followed by the *Proteobacteria* (22%), *Bacteroidetes* (14%), *Actinobacteria* (9%), *Chloroflexi* (2%), *Acidobacteria* (2%), Archaea (1%) and 34 other minor phyla. The 1% of Archaea is subdivided into *Euryarchaeota* (74%), *Crenarchaetoa* (13%), *Thaumarchaeota* (9%), *Woesearchaeota* (3%) and others (1%). Altogether, the DAIRYdb is able to capture the diversity of known taxa expected to occur in dairy products. Increasing number of whole genome sequences (WGS) will more likely lead to a replacement of incomplete 16S sequences in the DAIRYdb by full-length sequences that cover all HVRs.

**Fig. 3.**
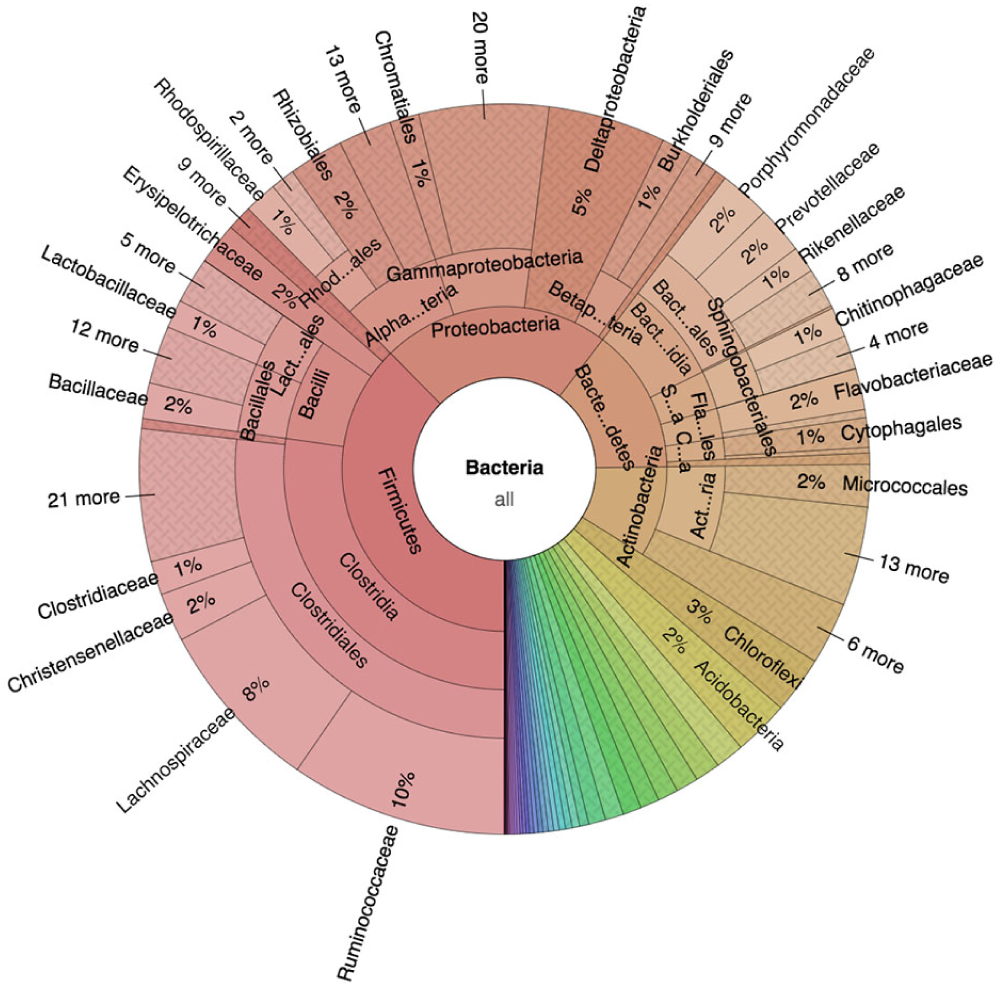
Complete microbial diversity present in the DAIRYdb. Prokaryotic biodiversity in the DAIRYdb is represented by 2 kingdoms, 47 phyla, 136 classes, 249 orders, 463 families, 1’757 genera and 4’030 unique species-like groups. The most represented phyla is Firmicutes (37% of all sequences), followed by the Proteobacteria (22%), Bacteroidetes (14%), Actinobacteria (9%), Chloroflexi (2%), Acidobacteria (2%), Archaea (1%) and 34 other minor phyla.

The cheese microbiome is often dominated by few phylogenetically closely related species, of lactic acid bacteria (LAB) belonging to a few genera (*e.g.*, *Lactobacillus*, *Lactococcus*, *Leuconostoc* and *Streptococcus*) (9). Therefore, special attention was put into the manual curation of the DAIRYdb sequences at species rank. Despite the genotypic and phenotypic characteristics of the most common LAB in cheese are extensively studied and described, several unresolved controversies regarding the nomenclature of some keystone species still remain unsolved, such as for the species *L. helveticus* and *L. gallinarum*, *S. thermophilus* and *S. salivarius*, *L. casei* and *L. paracasei* and *L. zeae*, *L. plantarum* and *L. paraplantarum* (77–79). The DAIRYdb is composed of sequences retrieved from the Silva database along with their respective taxonomy, which was manually inspected for nomenclature hierarchy conflicts based on the phylogenetic position within the tree. However, some conflicting annotations of the same sequence were detected between the Silva taxonomy and the Bacterial Diversity Metadatabase, such as the species assignment of the type strain sequence Accession AB008205, which is labelled as *L. casei* in Silva and *L. paracasei* in BacDive (80). For the reference sequences of the most crucial species, it was tried to use bacterial names listed in the actual "List of prokaryotic names" according to BacDive, however, further disagreements between Silva and BacDive cannot be completely excluded. Moreover, it is also possible that some crucial genera in dairy products may undergo a radical genome-based relabelling in order to have more homogeneous clusters (79).

Different approaches were applied on inpure taxa, *i.e.* taxa that overlap in the tree despite being assigned to different nomenclature (54), by the universal databases. For instances, for the genera *Escherichia* and *Shigella*, Silva, LTP and RDP use the combined genus name *Escherichia–Shigella* but retain well-established species names, such as *Escherichia coli*. Differently, Greengenes leaves their sequences unclassified at ranks below the family *Enterobacteriaceae* (54). The different taxonomic nomenclature references used by the three databases have an impact on revisions to resolve conflicts with sequence-based phylogenies and the labelling of new candidate groups identified in environmental sequences. However, discussion on the taxonomic inconsistencies and limitations of the universal databases (Silva, LTP, RDP and Greengenes), which the DAIRYdb was compared with, goes beyond the scope of this study and was extensively discussed elsewhere(54, 70).

The DAIRYdb will undergo regular updates in accordance to update on bacterial nomenclature (79), integrating the novelties or correcting the changes. Finally, the inclusion of full-length and high-quality 16S sequences from reference type strains leads to a more robust and confident taxonomic classification(68).

### Validation

At present, only short read sequences can be obtained from the most common amplicon NGS sequencer with at least 99% quality and up to 600 bp in length (Illumina MiSeq, Ion Torrent S5). Although long reads sequencing technology, such as PacBio and Oxford Nanopore, are steadily improving read quality, they are not yet routinely used for amplicon metabarcoding studies. Therefore, performance of the DAIRYdb was evaluated on short read sequences spanning over either a single HVR or HVR pairs. The single HVRs and HVR pairs were extracted from randomly subsampled sequences from the DAIRYdb using two methods: V-Xtractor (81) (Figure 4A,C) or *in silico* PCR with mothur (36). V-Xtractor was used to evaluate the general annotation accuracy of all HVRs present in the DAIRYdb. The *in silico* PCR with universal primers (Table 1) highlighted the theoretical extraction efficiency of the primer pairs adapted to the pool of sequences in the DAIRYdb. While V-Xtractor extracted the HVRs, the *in silico* PCR also evaluated the theoretical extraction efficiency of different primer pairs. The ratio between the number of detected HVRs with V-Xtractor and HVRs extracted by *in silico* PCR determined the biodiversity coverage of the different HVRs achieved with the different primer pairs and potential biases in community structure in downstream analyses depending on the different HVR analysed (Figure 4B,D).

**Table 1.**
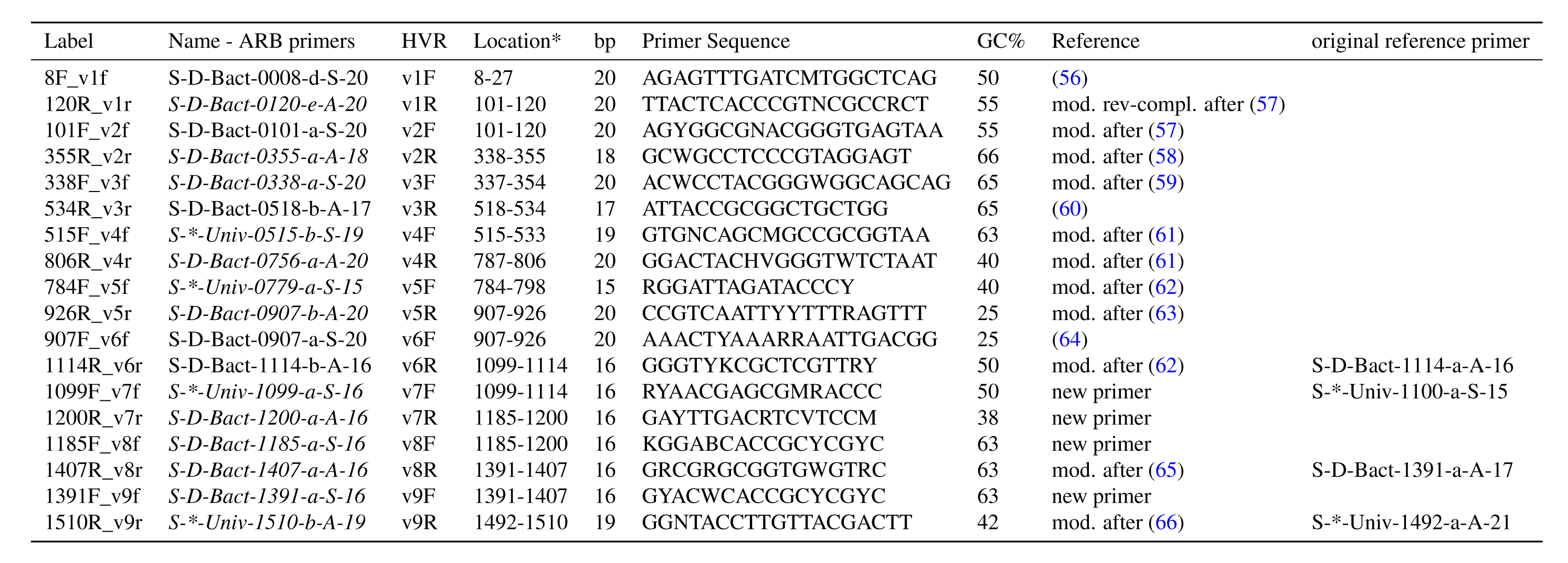
Primers used in the in silico PCR extraction of the HVRs. \**E. coli* position as a reference.

**Fig. 4.**
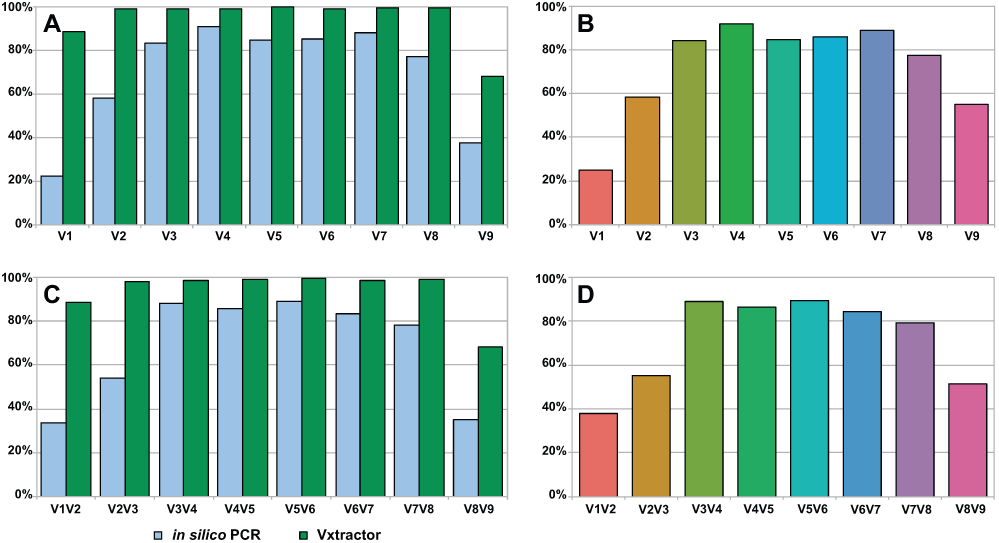
Presence and extraction efficiency of all HVR in the sequences of the DAIRYdb. Single HVR (A) and (B) and HVR pairs (C) and (D) HVRs were extracted using in silio PCR with mothur (B) and (D) and HVR extraction with V-Xtractor (A) and (B) from sequences present in DAIRYdb v1.1 to test completeness of the sequences therein. While almost 100% of the 10’290 sequences span over V2 to V8 only 89% contain V1 and 68% contain V9 (A) and (C). The in silico PCR highlights the theoretical amplification efficiency of the most common universal primers with 0 mismatches normalized to the total number of detected HVR (B) and (D).

Almost 100% of the sequences in the DAIRYdb span from V2 to V8. The HVR V1 (89%) and V9 (68%) are the regions with the least coverage in the DAIRYdb. This is due to the commonly used universal primers 8F and 1492R for the full-length 16S PCR leading to the entirely or partial loss of V1 and V9 (Figure 4A, Table 1). The primer pairs targeting the V1, V2 and V9 were less efficient as compared to the primer pairs targeting V3 to V8. The primer pairs for V4 performed best with 90% coverage, followed by V7 (88%), V5 and V6 (85%), and V3 (83%). The same hold true for the HVR pairs, where the HVR pairs V1V2, V2V3 and V8V9 performed less well as compared to the central HVR pairs (Figure 4C, 1).

The net *in silico* performance of each primer pair is presented as normalized to the total number of sequences detected by V-Xtractor for each HVR (Figure 4B,D). Percentage values of the single HVRs slightly increased while confirming the overall picture. The largest biodiversity coverage by the DAIRYdb was achieved by the single HVR V4 (92%), followed by the HVR pairs V5V6 and V3V4 (89%).

The taxonomy annotation accuracy of the DAIRYdb was compared with other universal databases, such as Silva128, RDP trainset v16, LTP and Greengenes analysing fragments of single HVRs or HVR pairs extracted from the sequences in the DAIRYdb with V-Xtractor and *in silico* PCR. The synthetic HVR fragments were extracted from 1000 subsamples of each 100 randomly selected sequences from the DAIRYdb by either V-Xtractor or *in silico* PCR and assigned to all taxonomic ranks by the means of three different classification predictors (Blast+, Metaxa2 and SINTAX) using the aforementioned databases. Taxonomic annotation accuracy of the single HVR extracted with V-Xtractor with the DAIRYdb using SINTAX was above 75% at all taxonomic ranks (Figure 5). Accuracy was highest for the even HVRs (V2, V4, V6 and V8) as compared to the odd HVRs (V1, V3, V5, V7 and V9). The region V2 presented the greatest classification accuracy, which is in line with other findings showing that the regions V1 and V2 resulted in a more accurate OTU clustering at 97%, 98% and 99% (32). Overall, the universal databases were less accurate with decreasing taxonomic rank (Figure 5B-D). Only the RDP trainset v16 achieved about 25% of correct species annotations, while the other databases only classified to genus rank. Although the RDP trainset v16 performed best among all universal databases, annotation accuracy was below the accuracy values assessed in previous studies (54). Different to the DAIRYdb, the HVR V4 performed best with the universal databases with exception to Silva, where V2 achieved a higher accuracy (Figure 5). Generally, the difference in classification accuracy was stable through all HVRs with exception to the Silva database, where bigger oscillations were observed between the HVRs showing a clear drop for V6 and V7 (Figure 5E). All HVRs taken together, the DAIRYdb achieved a significantly better taxonomy annotation accuracy of average 88.9% ± 5.5 as compared to the universal databases tested, particularly at order to species ranks (Figure 5F). The annotation accuracy results with Blast+ and Metaxa2 of single HVR extracted with *in silico* PCR (Additional File 3, Figures 1, 3 and 7), V-Xtractor (Additional File 3, Figures 5 and 9), and HVR pairs with *in silico* PCR (Additional File 3, Figures 2, 4 and 8), V-Xtractor (Additional File 3, Figures 6 and 10) are available in Additional File 3.

**Fig. 5.**
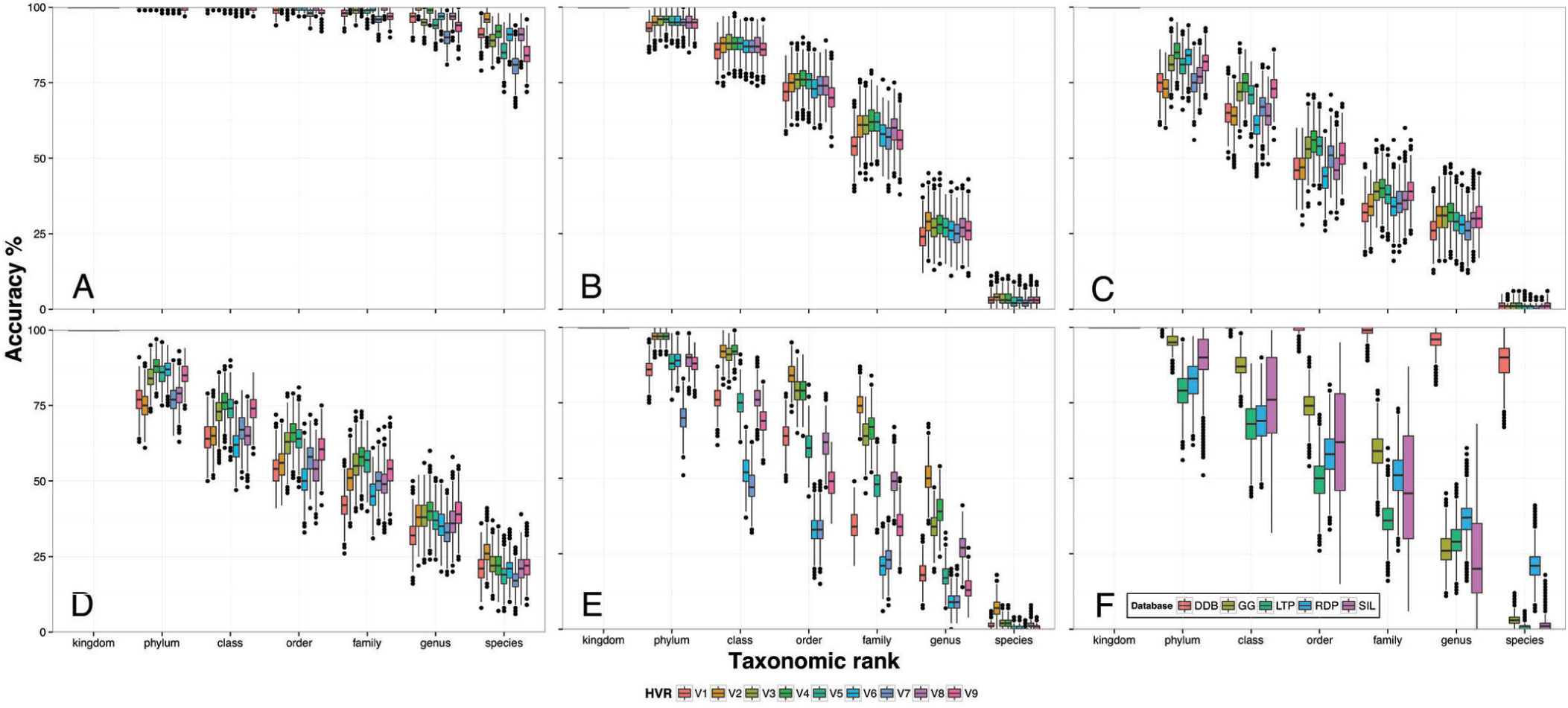
Taxonomy annotation accuracy of the DAIRYdb on reads extracted with V-Xtractor. The HVR pairs V1-V2, V2-V3, V3-V4, V4-V5, V5-V6, V6-V7, V7-V8, V8-V9 were re-annotated using three different classification algorithms, Blast+, Metaxa2 or SINTAX, respectively. This figure shows the results with SINTAX (Analyses with Metaxa2 and Blast+ are shown in Additional File 2). Taxonomy annotation was bootstrapped 1000 times with a subset of 100 randomly selected sequences from the DAIRYdb and annotated with DAIRYdb (A), Greengenes (B), LTP (C), RDP (D) and Silva (E). Average performance of all HVR for each database (F) (accuracy = correctly annotated/total).

The results with the HVR pairs was similar to the single HVRs (Figure 6). Classification confidence between HVR pairs was less variable between different HVR pairs and within the bootstrapping values of the same HVR pair as compared to the single HVRs, indication for a more robust classification with increasing number of HVRs. The HVR pair V1V2 achieved the highest classification accuracy at species rank in the DAIRYdb, as well as with RDP and Silva. These results are in agreement with previous studies, where V1 and V2 have been shown to have the highest average classification accuracy and average confidence estimate up to the genus rank (24). Greengenes species annotation accuracy was similar for all HVRs, while LTP showed very low performance at species rank. The average accuracy value for correct species annotation of all HVR pairs with the DAIRYdb was over 94% ± 2.8 (Figure 6F). Only species annotation with the RDP trainset v16 achieved 25% of correct annotations. The BLAST16S database was shown to obtain genus accuracies ∼50% for V4, which improves with increasing length to ∼60% with V3–V5 and ∼70% with full-length 16S (70). As expected, the increasing number of HVR increases the confidence in taxonomy annotation.

**Fig. 6.**
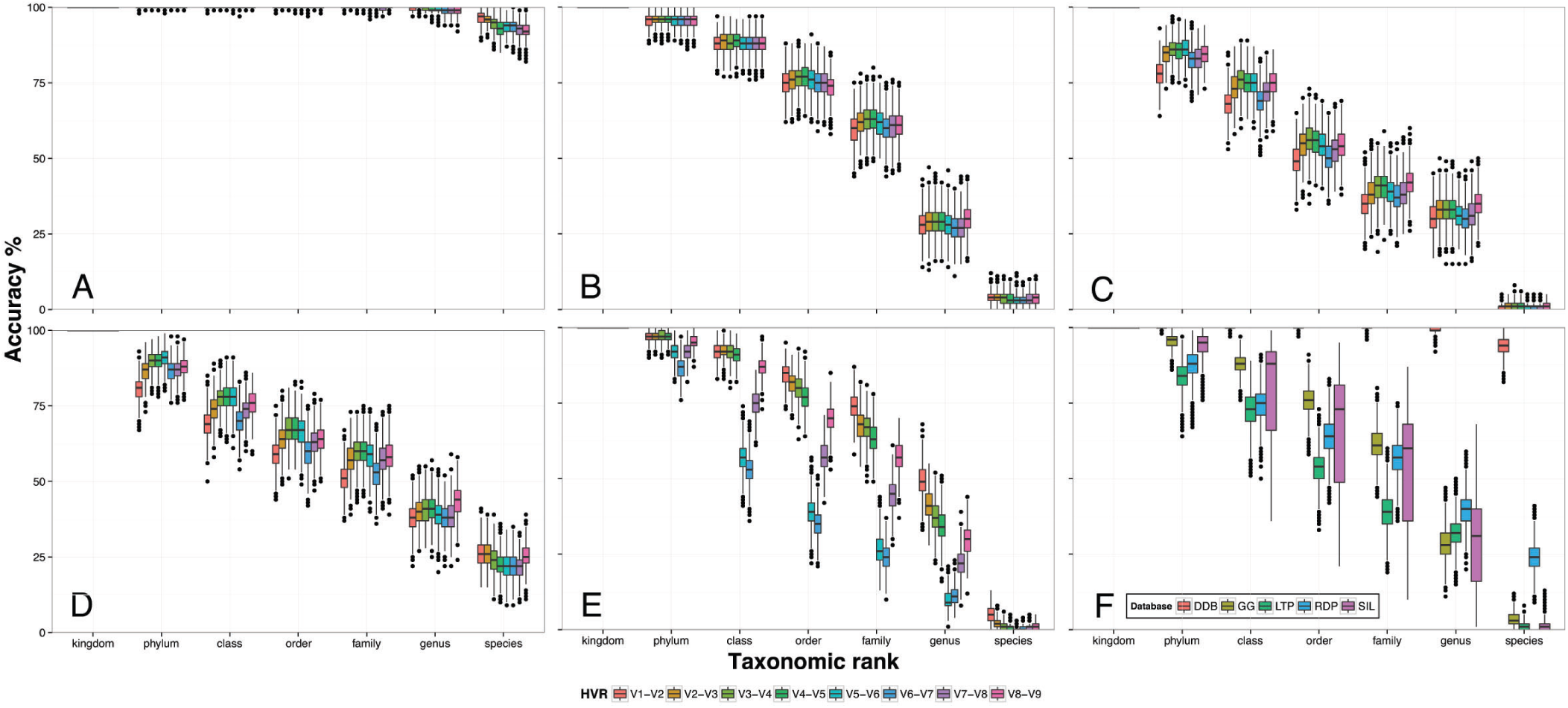
Taxonomy annotation accuracy of the DAIRYdb on reads extracted with V-Xtractor. The HVR pairs V1-V2, V2-V3, V3-V4, V4-V5, V5-V6, V6-V7, V7-V8, V8-V9 were re-annotated using three different classification algorithms, Blast+, Metaxa2 or SINTAX, respectively. This figure shows the results with SINTAX (Analyses with Metaxa2 and Blast+ are shown in Additional File 2). Taxonomy annotation was bootstrapped 1000 times with a subset of 100 randomly selected sequences from the DAIRYdb and annotated with DAIRYdb (A), Greengenes (B), LTP (C), RDP (D) and Silva (E). Average performance of all HVR for each database (F) (accuracy = correctly annotated/total).

Taxonomy annotation accuracy varied only little between different taxonomy predictors with the DAIRYdb and not significantly with either both, single HVR (Additional File 3, Figure 11) or HVR pairs and (Additional File 3, Figure 12). In fact, classification annotation accuracy performance varied more dependent on the database rather than the classification predictor. These results indicate that annotation of the members of the bacterial community is primarily influenced by the selection of the database, by the HVR, and only then by the taxonomy predictor (Additional File 3, Figures 1-10). A comparison of the three classification predictors, Blast+, Metaxa2 and SINTAX with the DAIRYdb confirmed that HVR pairs could be more accurately assigned to the correct species than single HVR (Figure 7). Among all tools, Blast+ and SINTAX were slightly yet not significantly better than Metaxa2. Since Metaxa2 uses more stringent parameters, as it only assigns the taxa if in agreement with Blast+, the lower performance of Metaxa2 with respect to Blast+ alone is not surprising. Moreover, Metaxa2 performance is strongly dependent on the average nucleotide identity (ANI) thresholds used, which were set according to (82). On the other hand, the more stringent parameters of Metaxa2 reduce the number of over-classified sequences. Generally, taxonomy annotation results are most robust whilst using different classification predictors with the DAIRYdb. We therefore recommend to use both, Metaxa2 with integrated Blast+ and SINTAX to obtain taxonomy annotations closest to the ground truth. Although a lower SINTAX cutoff of 0.6 increases the risk of over-classification, it is justified by the better quality of the DAIRYdb and the comparison with Metaxa2 for definitive taxonomy annotation (more details on the recommended usage on real samples are described on https://github.com/marcomeola/DAIRYdb).

**Fig. 7.**
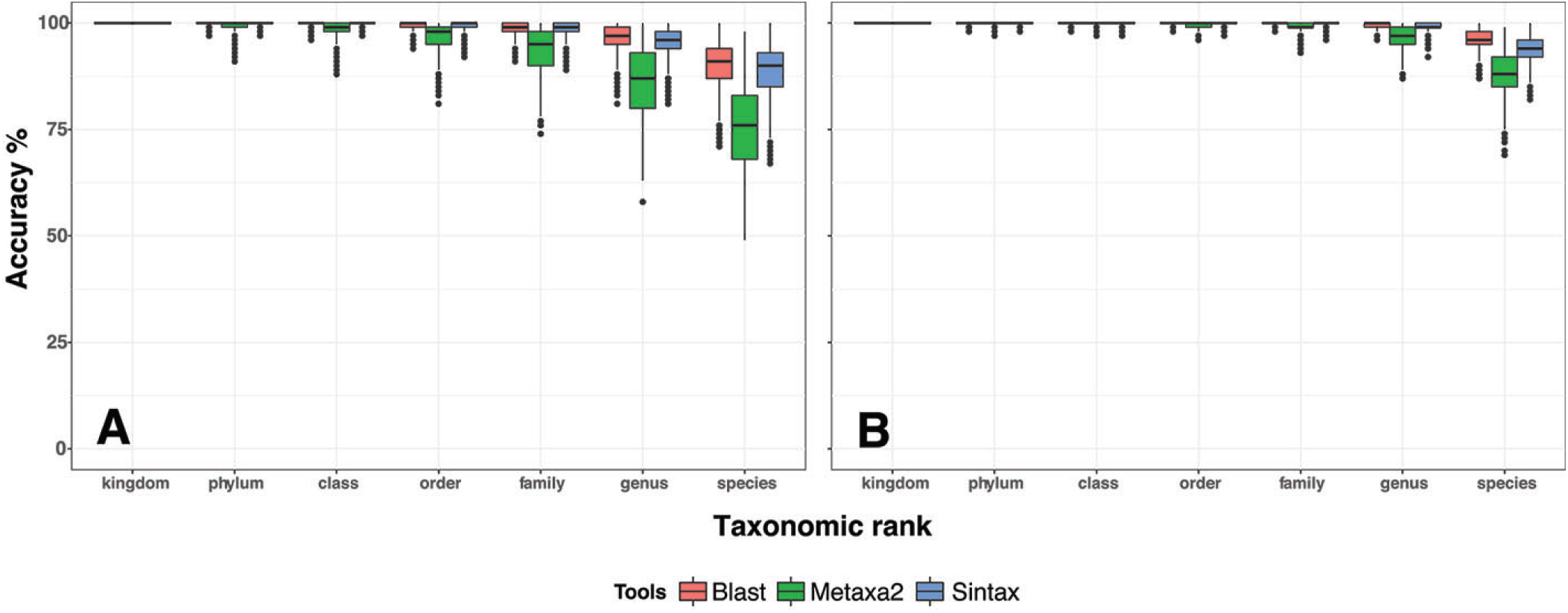
Comparison of the overall annotation accuracy of the three algorithms, Blast+, Metaxa2 and SINTAX for all single HVR (A) and HVR pairs (B). Although Blast+ presented a slightly better performance over SINTAX and Metaxa2, it was not statistically significant. All three classification tools assigned more than 75% of the sequences using the DAIRYdb as a reference for all HVR pairs.

### Outlook

The advances of genomics in microbiology has led to a reassessment of the phylogeny, which still remains a moving target particularly for microbial taxonomy (53, 54). The complexity of microbial taxonomy was already reflected in the statement by S. T. Cowan saying that "Taxonomy is the most subjective branch of any biological discipline, and in many ways is more an art than a science" (52, 83). As of 2014, the number of species of prokaryotes with validly published names was about 11’000 (84). However, microbial systematics is failing in its fundamental mission to precisely provide the ecological properties of an organism that is classified to a species (85).

The correct definition of a bacterial community structure remains a bioinformatic challenge, where any parameter, from wet-lab (*i.e.*, DNA extraction, primer and HVR selection, amplification, sequencing) to the bioinformatic pipeline, can influence the outcome. The results of microbiome studies are most strongly influenced by the selection of the primer pairs and thereof of the HVR amplified, rather than on the sequencing technology used for the study (86–88). The OTU-picking algorithm dependent on the sequencing technology (clustering vs. denoising) or ASVs instead (89), the classification predictor are of secondary importance, although their impact on the outcome is not negligible (67). The selection of the primer pairs should be made after careful consideration of their coverage in diversity with respect to the studied environment (67). Although researchers tend to use primers as universal as possible to catch the entire diversity present in the samples, it might be a pragmatic approach to lose some universality while increasing specificity for the studied environment. For dairy products, the DAIRYdb achieves both, covering all the biodiversity expected in these environments, while achieving specificity in taxonomic annotation.

The main scope of the DAIRYdb is to improve accurate species classification in dairy products. Beyond this, it covers a considerable diversity in agreement with the diversity detected in dairy products so far. However, the DAIRYdb does not necessarily perform better than universal databases on a set of sequences from another environments, such as the human gut. Classification accuracy performed on sequences from type strains included in the Human Intestinal Tract database (HITdb) showed that the DAIRYdb performed comparably well to the RDP trainset v16 and significantly better than Silva and Greengenes (Figure 8) (69). Yet, the way and ability to recognize the basic unit for taxonomy of prokaryotes depends on the resolution power of the observational methods actually available (52). The study of every particular environment calls upon peculiar requirements. Dairy products are no exception, as their bacterial communities are usually dominated by few phylogenetically highly related species, which are often difficult to discern, such as *L. casei*, *L. paracasei* and *L. rhamnosus* or *S. thermophilus* and *S. salivarius*. Particularly for *S. thermophilus*, which is a very important representative bacterium in dairy products, the official name still is *S. salivarius subsp. thermophilus* (90). Although a separate full species status was proposed (91), persistent contention prevented a full ratification by the taxonomic committees (90). Increasing sequence read lengths will make it possible to cover three HVRs or even the entire 16S, thus significantly improving taxonomic annotation accuracy at species rank by using a manually curated database like the DAIRYdb.

**Fig. 8.**
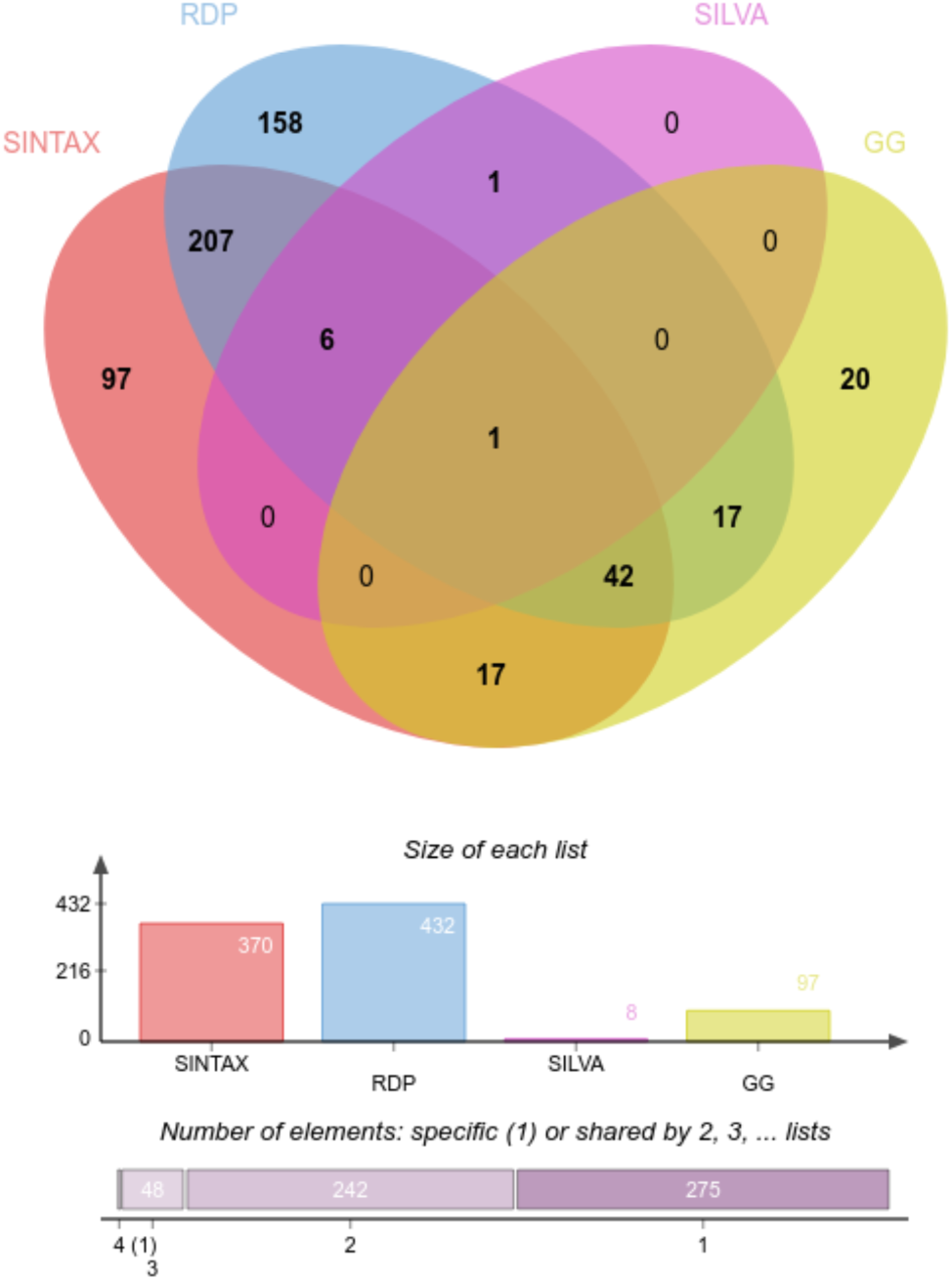
Taxonomy annotation accuracy test on sequences from the HITdb. Comparison of the taxonomy annotation accuracy at species rank between the DAIRYdb, RDP, Silva and Greengenes on type strain sequences present in the HITdb (69). On sequences from origin other than dairy products, the DAIRYdb performs significantly better than Silva and Greengenes, but not better than the RPD trainset v16.

Although it can be considered a significant progress to obtain over ∼90% of accurate species classification based on short 16S fragments, quality of dairy products is often influenced by different strains of the same species. The resolution at strain or subspecies rank, however, based on full 16S is highly unlikely to be achieved independently from advancing sequencing technology. While on the one hand the definition of strains and subspecies is even more problematic than higher ranks such as species (85), on the other hand, the intraspecies variability of the 16S lacks sufficient resolution to clearly discern between strains and subspecies within the same species (92). Nevertheless, recent powerful bioinformatics tools, such as Oligotyping (93) or Minimal Entropy Decomposition (MED) (94), can be applied to distinguish between ecologically relevant amplicon sequence variants (ASVs) within OTUs assigned to a same species. The resulting oligotypes or haplotypes within a species might be linked to different metabolic pathways or associated to identified physico-chemical characteristics of cheese or dairy products. Hereof, the DAIRYdb is a powerful improvement as it accurately identifies the sequences belonging to the same species, which can further be decomposed to oligotypes. Finally, links between oligotypes and 16S from WGS could improve the link between phylogeny and ecotypes for a better ecological understanding of the system (85, 89, 95).

## Conclusions

Accurate prediction of taxonomy based on the marker gene 16S is a fundamental step in microbial diagnostics and microbial ecology studies. Dairy products, particularly cheeses, are enriched by a few dominant species often belonging to the same genera, such as *Lactobacillus spp.*, *Lactococcus spp.*, *Streptococcus spp.*. An automatic and reliable taxonomic annotation to the correct species is pivotal to further routine microbial diagnostics.

While universal 16S databases, such as Silva, RDP and Greengenes cover a broad biodiversity allowing to capture the maximal biodiversity available in a system, the enormous number of sequences in those databases lead to conflicting taxonomy annotation at genus and species ranks and ambiguous annotations or blanks due to competing sequences increase accordingly (69). Moreover, the size of the database can be a deterrent for researchers to improve the quality of taxonomic annotation of the sequences therein. Most of the detected OTUs in NGS analyses diverge from authoritative reference sequences from type strains either due to sequencing biases or missing cultivated representative strains. Beside reference sequences, many environmental sequences are annotated by the universal databases Silva, RDP and Greengenes based on different taxonomic classification standards, e.g., Bergey’s Manual, the List of Prokaryotic Names with Standing Nomenclature (LPSN), International Nucleotide Sequence Database Collaboration (INSDC). These different curation strategies lead to annotation conflicts between the databases and disagreement between microbiome studies, which are not biologically explained, rather a consequence of the database used for annotation.

Different to available universal databases, DAIRYdb achieved correct taxonomy annotation for ~90% of species names on single HVRs and HVR pairs with sequences present in dairy samples (70). In fact, the DAIRYdb significantly reduced conflicting miss-annotated sequences and facilitated manual curation, while covering the inspected biodiversity. The better performance of the DAIRYdb over universal databases can be explained by the overall reduced number of sequences, only 10’290, with no conflicting taxonomy at all taxonomic ranks. Our results are in disagreement to the recommendation to use the largest and most diverse database possible for 16S classification (96). On the opposite, manually curated 16S databases with authoritative fulllength 16S sequences dedicated to the studied environment enormously improve classification confidence to the species rank (54, 68, 69). Reducing the number of representative sequences to a minimal number in the training set further diminishes the risk of highly similar sequences with conflicting taxonomy, thus lowering the performance of the database used for classification (54, 68).

A certainly valid argument against manually curated databases is their lack of reproducibility (54). However, annotation accuracy achieved with the DAIRYdb significantly outperformed all universal databases tested here, as well as the RDP trainset v16, which was shown to have the best performance among the universal databases (54). The training sets of the universal databases contained sequences with missing taxonomic labels at uncertain classifications at any taxonomic rank with increasing number of blanks at species levels (54). The consequences are numerous unclassified OTUs with no biological meaning.

We therefore propose the manually curated DAIRYdb as a gold standard database for 16S microbiome studies on cheese and dairy products. The implementation of a curated database may lead to wider consensus and standardization processes reducing conflicts in literature due to the use of different universal databases integrated in different classification tools (67, 97).

## ACKNOWLEDGEMENTS

Eric Dugat-Bony (INRA Grignon, GMPA) improved the completeness of the DAIRYdb by providing some additional 16S sequences from his own studies related to cheese rind bacteria. We are grateful to the genotoul bioinformatics platform Toulouse Midi-Pyrenees (Bioinfo Genotoul) and INRA MIGALE bioinformatics platform (http://migale.jouy.inra.fr) for providing computing and storage resources.

